# Frontal midline theta power during the cue-target-interval reflects increased cognitive effort in rewarded task-switching

**DOI:** 10.1101/2023.09.25.559275

**Authors:** Stefan Arnau, Nathalie Liegel, Edmund Wascher

**Author notes:** **Corresponding author:** Stefan Arnau, Leibniz Research Centre for Working Environment and Human Factors Dortmund (IfADo), Ardeystraße 67, 44139 Dortmund, Germany.

## Abstract

Cognitive performance largely depends on how much effort is invested during task-execution. This also means that we rarely perform as good as we could. Cognitive effort is adjusted to the expected outcome of performance, meaning that it is driven by motivation. The results from recent studies investigating the effects manipulations of motivation clearly suggest that it is the expenditure of cognitive control that is particularly prone to being affected by modulations of cognitive effort. Although recent EEG studies investigated the neural underpinnings of the interaction of effort and control, reports on how cognitive effort is reflected by oscillatory activity of the EEG are quite sparse. It is the goal of the present study to bridge this gap by performing an exploratory analysis of high-density EEG data from a switching-task using manipulations of monetary incentives. A beamformer approach is used to localize the sensor-level effects in source-space. The results indicate that the manipulation of cognitive effort was successful. The participants reported significantly higher motivation and cognitive effort in high versus low reward trials. Performance was also significantly increased. The analysis of the EEG data revealed that the increase of cognitive effort was reflected by an increased mid-frontal theta activity during the cue-target interval, suggesting an increased use of proactive control. Alpha-desynchronization throughout the trial was also more pronounced in high reward trials, signaling a bias of attention towards the processing of external stimuli. Source reconstruction suggests that these effects are located in areas related to cognitive control, and visual processing.

## Introduction

Performance in a cognitive task not only depends on skill but is also crucially dependent on how much effort is invested during task-execution. Shenhav and colleagues (2017) defined mental effort as the set of intervening processes that connect task characteristics and the individual capacity of the performing individual on the one side, and the fidelity of the information-processing operations on the other side. This means that effort translates skill to performance. On the flipside, it also means that effort is usually limited, and we rarely perform as good as we could. The reason for this limitation is that, apart from specific phenomena, like flow experience (van der Linden et al., 2020), or learned industriousness (Eisenberger, 1992; Inzlicht et al., 2018), cognitive effort feels aversive, and humans usually tend to avoid it (Kurzban, 2016). The subjective experience of effort can thus be seen as an integral part of a resource management system that adjusts the level of task engagement to the expected outcome of successful performance (Shenhav et al., 2013). The investment of cognitive effort is consequently driven by motivation.

In cognitive psychology, this dependency of cognitive effort on motivation has often been investigated using the monetary incentive delay paradigm (Knutson et al., 2000). Numerous studies that deployed this paradigm demonstrated that motivation and mental effort are linked. The strongest effects of reward-related modulations of effort could be observed for tasks requiring cognitive control, like the Stroop task (Padmala & Pessoa, 2011), the stop-signal task (Boehler et al., 2014), task-switching (Kleinsorge & Rinkenauer, 2012; Shen & Chun, 2011), a working memory task (Heitz et al., 2008), and visual search (Navalpakkam et al., 2009). Cognitive control is a collective term for a set of superordinate functions that are responsible for encoding and maintaining task representations, as well as for coordinating cognitive functions that are necessary for the execution of a task (Botvinick & Braver, 2015). The observation of cognitive control being affected by motivation makes sense from a resource-management perspective, as cognitive control constitutes a bottleneck in human information processing (Feng et al., 2014; Shenhav et al., 2017). Reasons for these limitations of processing capacity may be constraints in working memory capacity (Cowan, 2001; Cowan et al., 2012; Oberauer et al., 2016), or interference due to concurrent cognitive processes requiring the same resource at the same time (Allport et al., 1972; Wickens, 2008).

Neuroscientific methods have been deployed to investigate how cognitive effort is adjusted to different reward contingencies. Studies using fMRI demonstrated that the availability of reward is linked to a stronger activation in dopaminergic areas like the ventral striatum (Cubillo et al., 2019), but also to activation in frontal and parietal areas that are associated with cognitive control (Egner, 2017; Jimura et al., 2010; Krebs et al., 2012; Padmala & Pessoa, 2011). It has also been observed that the dorsal anterior cingulate cortex (dACC) is involved in reward-based decision making (Wallis & Kennerley, 2011). Recent theories assume that the dACC integrates reward-related information to the exertion of cognitive control (Shenhav et al., 2013), subsequently affecting attentional engagement, strategy selection, and control-intensity (Chong et al., 2017; Shenhav et al., 2016). A recent meta-analysis investigated aggregated findings of fMRI studies investigating motivated cognitive control and identified the mid-dorsolateral prefrontal cortex, the anterior insula, the intraparietal sulcus, as well as the anterior mid-cingulate cortex being consistently activated in reward versus no reward trials (Parro et al., 2018).

An important aspect when investigating cognitive effort using neuroscientific methods is to consider which cognitive processes are present in the obtained signal. In fMRI as well as in EEG studies, the signal is usually analyzed time-locked to experimental stimulation. Depending on the experimental design used, these processes may vary substantially. In rewarded task-switching, some studies presented the reward-cue and the task-cue at the same time, but prior to the target stimulus (e.g. Shen & Chun, 2011), while some studies present reward-cues, task-cues, and targets one after the other (e.g. Hall-McMaster et al., 2019). Due to its high temporal resolution, the EEG can disentangle effects of cognitive effort on cognitive processes if reward-cues, task-cues, and target stimuli are presented sequentially. Neural signals in response to reward-cues seem to reflect an automatic appraisal process and not just the resulting adjustment of cognitive effort (c.f. Frömer et al., 2019). In contrast, the analysis of physiological signals that are time-locked to the task-cue or the target stimuli allows for the investigation of preparatory processes and target processing.

Regarding target processing, recent studies could show that an increase of cognitive effort leads to a modulation of the intensity of cognitive engagement. In high compared to low reward conditions, a larger parietal P3 amplitude was observed (Schevernels et al., 2014; van den Berg et al., 2014), which indicates an enhanced allocation of cognitive resources (Kok, 2001). Van den Berg and colleagues (2014) found a more pronounced frontal P2 and posterior N2 component in high versus low reward trials, indicating a strengthening of attentional engagement. Also, a stronger N2 posterior contralateral (N2pc) was observed, indicating the selective orienting of visual attention in response to a lateralized stimulus presentation (Hickey et al., 2010; Sawaki et al., 2015).

When investigating cognitive effort, reward-related modulations of electrophysiological activity in response to the task-cue are probably even more relevant than target-related activity: First, preparatory processes during the cue-target interval require top-down cognitive control for updating or maintaining the task-rules, the response-mappings, and for directing attention. Effects of motivation on cognitive control can thus be observed more process-pure compared to the period after the imperative stimulus, when the task is executed. Second, modulations of preparatory processes may also indicate reward-related changes on the strategic level of task processing. Studies using the AX Continuous Performance Task (Servan-Schreiber et al., 1996) demonstrated a shift from a reactive to a proactive control mode in the prospect of reward (Chiew & Braver, 2014; Fröber & Dreisbach, 2016). According to the dual-mechanisms-of-control framework, a reactive control mode allocates cognitive control to a task when needed, for example in response to a target stimulus, or as a late correction mechanism. The proactive control mode, in contrast, is characterized by a constant allocation of resources, as action goals and task representations are actively maintained. It requires more effort but biases the cognitive system towards effective task-processing (Braver, 2012; Braver et al., 2008).

EEG studies showed that, in high versus low reward conditions, a stronger alpha desynchronization could be observed during the cue-target interval in a visual search task (Sawaki et al., 2015), as well as in a Stroop task (van den Berg et al., 2014). It has also been observed that the amplitude of the Contingent Negative Variation (CNV; Walter et al., 1964) is increased in high reward conditions during the cue target interval (Frömer et al., 2021; Schevernels et al., 2014; van den Berg et al., 2014), indicating a strengthening of preparatory attentional processes (Kononowicz, 2016). Hall-McMaster and colleagues (2019) used a representational similarity analysis on EEG data recorded in a task-switch paradigm. They demonstrated that the representation of the task rule during the cue-target interval is increased by reward, which also suggests that reward increases proactive control.

Although, as reported above, several recent studies investigated reward-related modulations of cognitive effort using the EEG, the literature regarding effects on oscillatory activity is quite sparse (c.f. Meyer et al., 2021). Apart from Sawaki and colleagues (2015), who performed an exploratory analysis in time-frequency space and observed a reduced alpha desynchronization during the cue-target interval of a switching-task, most studies analyzed specific measures to test their hypotheses (e.g. Hughes et al., 2013; Kang et al., 2018; van den Berg et al., 2014). Therefore, the overall picture of how motivation-related modulations of cognitive control are reflected in the time-frequency decomposed EEG signal is still incomplete. To bridge this gap, the present study aims at performing an exploratory analysis of high-density EEG data from rewarded task-switching. As no recent EEG study reports reconstructed sources for cognitive effort, a beamformer approach will be used to locate the observed effects in source space by using forward-models computed from individual structural MRI scans. This might enable us to connect time-frequency resolved effort-related oscillatory activity to the findings from recent fMRI studies.

## Method

### Sample

The participants were recruited via an announcement in a Facebook group of the Leibniz Research Centre for Working Environment and Human Factors Dortmund. All participants were screened for color blindness (Ishihara and Yoshii, 1972), reported normal or corrected to normal vision, and stated not to suffer from any psychiatric or neurologic disorders. It was also ensured that the participants met the criteria for the acquirement of a structural MRI scan. Overall, the data of 27 participants were acquired in this study. One dataset had to be excluded due to missing event-markers in the EEG data. The remaining sample of *N* = 26 (23 female) had a mean age of *M* = 21.65 years (*SD* = 2.15). For the source-space analysis, three additional datasets had to be excluded due to erroneous data from the anatomical MRI scan or the 3d scan of the head shape. All participants signed an informed consent before participating in the experiment. The study was approved by the ethics committee of the Leibniz Research Centre for Working Environment and Human Factors, Dortmund, and was in accordance with the Declaration of Helsinki.

### Procedure

To control for circadian differences, all participants arrived at the institute at 8:30 a.m. in the morning. After conducting a COVID 19 rapid test and the preparation of the EEG cap, the participants received a pre-experimental briefing regarding the monetary compensation consisting of a fixed amount of 35 € and a variable amount of up to 20 €. They were instructed to respond as quickly and as accurately as possible to maximize the obtained score and the variable amount of the compensation.

The participants performed in a switching task adopted from Hubbard and colleagues (2019). A schematic depiction of the experiment is presented in Figure 1. The task was presented on a 20” color monitor with a resolution of 1024×768 pixels and a refresh rate of 100 Hz which was located at 1.4 meters distance to the participants. All colors are reported in CIE 1931 XYZ color-space values. If not reported differently, all stimuli were presented in black color (0.287 0.312 0.0) against a light grey background (0.287 0.312 12.5). The presentation of the switching task occurred in alternating sequences of eight trials of low and high reward. For a response that was correct and fast enough, the participants could earn one point in low reward trials, and ten points in high reward trials. The sequences began with the presentation of a reward cue (“---” for low and “$$$” for high reward) for 800 ms. Throughout the sequences, the current reward condition was indicated at any time by a colored frame at the edge of the screen in either yellow (0.36 0.43 50.0) or blue (0.222 0.282 50.0) for the low and high reward condition, counterbalanced across participants. At the end of each sequence, a feedback screen indicated the earned points at a central position of the screen for 800 ms. Within these sequences, each trial started with the presentation of a task-cue, consisting of the letter A, B, X, or Y (1° viewing angle height) for 200 ms followed by a cue-target interval displaying a fixation cross for 800 ms. Subsequently, a probe display consisting of a circular arrangement (6° diameter relative to the center of the Gabor patches) of eight Gabor-patches (1.5° diameter each) was presented on the screen until a response was given or until a maximum response time of 1200 ms has passed. Depending on the task cue, the participants had to either perform a color-discrimination task by responding to the Gabor-patch with a deviating color (either red [0.5 0.38 25.0] or green [0.264 0.456 25.0] at peak saturation), or to perform an orientation-discrimination task by responding to the Gabor-patch with a deviating orientation (tilted left or right by 20°). The task-cues A and B were mapped to one of these tasks, the task-cues X and Y to the other. This mapping was counterbalanced across participants. For the color discrimination task, the mapping of green and red to the left and right response keys was also counterbalanced across participants, whereas it was fixed to a corresponding mapping for the orientation task (left tilt to left response, right tilt to right response) to avoid Simon effects (c.f. Simon, 1969).

**Figure 1:**
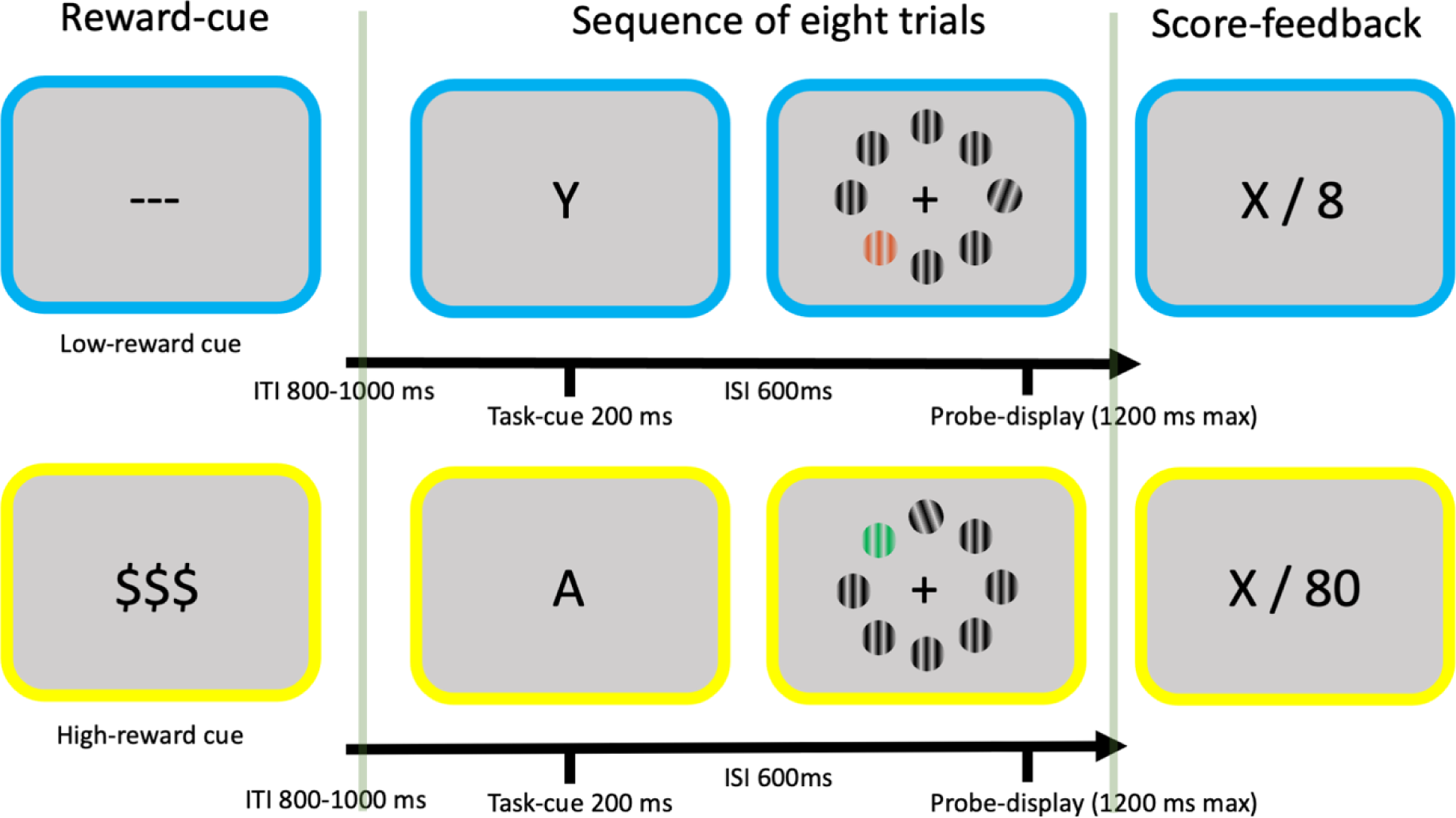
This figure shows a schematic depiction of a sequence of trials. Each sequence consists of a reward-cue indicating whether a low or a high reward is available for the upcoming trials, a sequence of eight trials, and a subsequent feedback screen informing about the awarded points (“X” refers to the respective points, “ITI” refers to “inter-trial-interval”, and “ISI” refers to “inter-stimulus-interval”). The task-cue indicates whether the task in the respective trial consists in a color-discrimination or an orientation-discrimination.

The experiment started with a series of four practice blocks. In the first two practice blocks, the participants learned the response mapping for the color and the tilt task in 16 trials, respectively. The next block of 32 trials presented both tasks as a switching task. The fourth practice block of 64 trials introduced the sequences of low and high reward. The 60^th^ percentile of the response times observed in this block served as the threshold for a response being classified throughout the experiment. The practice blocks were followed by eight experimental blocks with 256 trials each. After the experiment, the participants gave subjective ratings about their task-experience in a questionnaire consisting of analogue Likert scales. The participants rated the extent to which they paid attention to the reward condition at all. Separately for the low and the high reward condition, they also reported how motivated they were to show a good performance, how much effort they put into the task, and the extent to which they experienced mind wandering.

### Analysis of behavioral data and subjective ratings

For response time (only correct trials) and response accuracy repeated measures ANOVAs were calculated using a 2 × 2-design with the factors reward-condition and task-switch. After the experiment, the participants reported their subjective levels of effort, motivation, and experienced mind-wandering for low and high reward trials on analogue Likert scales. For all three measures, t-tests were calculated to test for effects of the factor reward-condition. For all analyses, effect sizes were calculated as bias-corrected partial η^2^ (Mordkoff, 2019; subsequently referred to as η^2^). Effect sizes are classified and discussed as small, medium, or large according to conventions of Cohen (1992).

### EEG recording and preprocessing

The EEG was recorded using 128 passive Ag/AgCl electrodes (Easycap GmbH, Herrsching, Germany) with a NeurOne Tesla AC-amplifier (Bittium Biosignals Ltd, Kuopio, Finland). During recording, a 250 Hz low-pass filter was applied. The ground electrode was at the position AFz, and the reference electrode was placed at FCz. Electrode impedances were kept below 10kΩ.

The preprocessing of the EEG data was performed using MATLAB (The Math Works Inc., Natick, Massachusetts) with the open-source toolbox EEGLab (Delorme & Makeig, 2004). The first step was to re-reference the data to the CPz electrode to obtain the signal for the FCz electrode. After resampling the data at 200 Hz, larger noise artifacts were removed from the signal by cutting out one-second-long segments after boundary events, and by rejecting continuous portions of the data based on spectral thresholding in the frequency range from 20 to 40 Hz, addressing muscular artifacts. The data were then band-pass filtered from 1 to 40 Hz and noisy channels were removed based on kurtosis criteria. On average, *M* = 8.23 (*SD* = 4.05) channels were rejected. Subsequently, the data were re-referenced to common average reference and segmented into epochs ranging from -1000 to 2600 ms relative to the onset of the task-cue. Noisy epochs were detected and removed automatically, with *M* = 226.69 (*SD* = 85.52) removed epochs on average. Finally, an independent components analysis was computed after compressing the data to the dimensionality matching its rank using PCA. In independent component (IC) space, ICs representing artifacts were identified using the ICLabel classification tool (Pion-Tonachini et al., 2019). ICs were regarded as artifacts and removed from the data if the classification results showed either a probability of less than 0.3 for the brain category, or a probability of more than 0.3 for the eyes category. On average, *M* = 70.04 ICs (*SD* = 10.97) were removed.

### Time frequency decomposition and analysis

The time frequency analysis was performed by convolving the EEG data with complex Morlet-wavelets. Trials from practice blocks as well as the first trials of each sequence were excluded from the analysis. The reason for excluding the latter is that an assignment to either the repetition or the switch condition was not possible. Twenty wavelets with logarithmically spaced frequencies ranging from 2 to 20 Hz were used. The widths of the tapering Gaussians were selected so that the corresponding full-widths at half-maximum of the Morlet-wavelets ranged from 1.25 to 2.75 Hz in the frequency domain and from 600 to 300 ms in the temporal domain (M. X. Cohen, 2019). To remove edge artifacts, the segments were pruned to -500 to 2000 ms relative to the stimulus onset of the task-cue. To perform the group-level analysis, the data were decibel-normalized relative to a frequency-specific baseline ranging from -500 to -200 ms before the onset of the task-cue. The baseline was calculated based on the average of the trials of all conditions (c.f. Arnau et al., 2020). This is important, as systematic differences in the baseline can be expected due to the sequences of low and high reward trials. In contrast to applying a baseline normalization for individual conditions, such systematic variance in the baseline period can still be observed.

For the group-level analysis of the electrophysiological measures in time × frequency × sensor space, cluster-based permutation tests were performed (Maris & Oostenveld, 2007). Cluster-permutation tests control for Type I error rates, which is essential when working with multiple comparisons in high-dimensional data. For each data point, *t*-statistics are computed, and a clustering algorithm identifies clusters of neighboring data points that are associated with a *t*-value corresponding to *p* < 0.1. The test statistic for each cluster is then defined as the sum of the *t-*values of all data-points within the cluster. This test statistic is evaluated against a H0-distribution obtained from collecting the maximum test statistics determined in 1000 iterations of a randomization procedure. In a two-sided test, clusters with *p* < 0.05 were regarded as significant. To test for effects of the factor reward, the assignment of the individual averages to low and high reward was permuted in the randomization procedure. To test for effects of the factor task-switch, the randomization procedure utilized the averages of repeat and switch trials. Interaction effects were tested by computing the differences of repeat and switch trials for low and high reward sequences separately and then test these differences as described for the main effect analyses. Again, effect sizes are reported as partial η^2^ (Mordkoff, 2019).

### Structural MRI acquisition and source-reconstruction

The goal of the source-level analysis was to reconstruct the cortical sources of oscillatory activity for frequencies and time windows for which significant effects could be observed at sensor level. To this end, individual anatomical MRI images on a Siemens 3T Prisma scanner with a 64-channel head/neck coil were acquired. The scanner was located at IfADo and T1-weighted data were obtained with the following parameters: MP-RAGE; TR = 2530 ms; TE = 2.36 ms; flip angle = 7°; 256 slices; matrix size = 256 × 256; resolution = 1 mm × 1 mm × 1 mm; acquisition time = 6 minutes and 30 seconds. To obtain digitized electrode positions and head shapes for each participant, a 3D scanning device was used (Structure Sensor, Occipital Inc., Boulder, Colorado). The MATLAB toolbox fieldtrip was used for all analysis steps (Oostenveld et al., 2011). First, the Structure sensor output and the T1-weighted MRI scan were aligned, and the individual electrode positions were manually derived from the 3D scan. Then, boundary element models with three layers (scalp, skull, and brain) were computed, based on the individual, aligned T1 images using the ‘dipoli’ method (Fuchs et al., 2002; Oostendorp & Van Oosterom, 1991). Each participant’s brain volume was then subdivided into a regular grid with 5 mm resolution. Grid points were warped as implemented in the fieldtrip function ‘ft_prepare_sourcemodel‘, so that each location fits a homogenous location in a 5 mm template grid. The template grid was derived from the MNI152 template brain used in fieldtrip (Holmes et al., 1998). The warping of the grid made it possible to compare the source estimates across subjects, and to plot the averaged source estimates on MNI152 template anatomical data.

For each grid point, we computed a common beamformer filter using all trials irrespective of the experimental condition. To this end, a leadfield matrix was computed for each grid point and the preprocessed EEG data was subdivided into epochs ranging from -800 to 2200 ms relative to the task-cue onset. Based on this data, cross-spectral density matrices were computed for each source-level comparison at the respective frequency specified above (see Result-section). A DICS Dynamic Imaging of Coherent Sources (DICS) beamformer was then derived from the cross-spectral density matrices and the leadfields (Gross et al., 2001).

In the second part of the analysis, the derived spatial filters were applied to EEG data of the experimental conditions. Short time Fourier transformations with multiple tapers were calculated for the preprocessed data for the relevant experimental conditions using the respective time windows and frequency (see Result-section). Then, the beamformer was applied to each power estimate. To correct for the center-of-head-bias of beamforming, the activation at each grid point was normalized using its noise estimate (c.f. Westner et al., 2022). The derived activation values for each source location were then used for statistical analysis. A cluster-based permutation test was conducted for each source level comparison. Dependent sample *t*-tests on grid point’s activation estimates were run. A clustering threshold for the grid points of *t*-values corresponding to *p* < 0.05 was used and the sum of t-values was used as test statistic. In a two-sided test, clusters with *p* < 0.05 were regarded as significant. The H0-distribution was derived from Monte Carlo simulations with 4000 runs. Effect sizes are reported as partial η^2^ (Mordkoff, 2019) for voxels belonging to significant clusters.

## Results

### Behavioral data and subjective ratings

The behavioral data as well as the subjective ratings assessed after the experiment are depicted in Figure 2. Analysis of the response times using repeated measures ANOVA showed a significant main effect for the factor reward (*F*(25, 1) = 21.51, *p* < .001, η^2^ = 0.45). Responses were faster in high compared to low reward trials (*M* = 495.01 ms, *SD* = 87.39 for low reward vs. *M* = 481.74, *SD* = 84.32 for high reward). Response times also differed significantly regarding the factor task-switching (*F*(25, 1) = 51.5, *p* < .001, η^2^ = 0.67), with faster responses for repeat compared to switch trials (*M* = 475.11 ms, *SD* = 80.56 for repeat vs. *M* = 501.65 ms, *SD* = 91.26 for switch). The interaction effect was not significant (*F*(25, 1) = 1.62, *p* = .214, η^2^ = 0.02).

**Figure 2:**
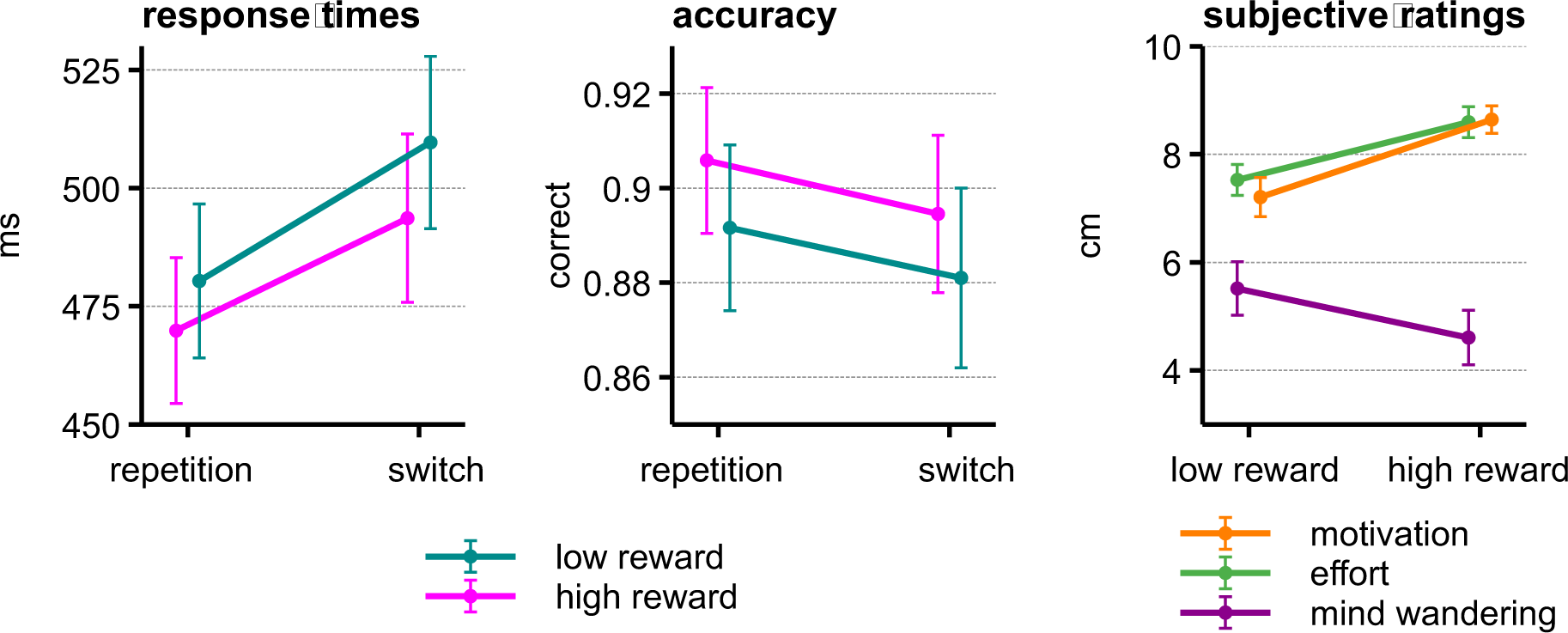
This figure depicts the behavioral measures response times and response accuracy as well as the subjective ratings for motivation, effort, and mind wandering. Error bars are depicted as standard error of means.

For response accuracy, the main effect for the factor reward was not significant (*F*(25, 1) = 3.78, *p* = .063, η^2^ = 0.1). The accuracy was *M* = 0.89 (*SD* = 0.09) for low reward and *M* = 0.9, (*SD* = 0.08) for high reward. Response accuracy differed regarding the factor task-switching (*F*(25, 1) = 6.58, *p* = .017, η^2^ = 0.18). Responses were more accurate in repeat compared to switch trials (*M* = 0.9, *SD* = 0.08 for repeat vs. *M* = 0.89, *SD* = 0.09 for switch). Again, the interaction effect was not significant (*F*(25, 1) = 0.01, *p* = .93, η^2^ = -0.04).

After the experiment, the participants rated their level of motivation significantly lower for low reward compared to high reward trials (*t*(25) = -4.3, p < .001, η^2^ = 0.117). The values (in cm) on the analogue Likert scales were *M* = 7.21 cm (SD = 0.36) for low reward and *M* = 8.64 cm (SD = 0.25) for high reward trials. Participants also reported significantly lower effort for low compared to high reward trials (*t*(25) = -4.53, p < .001, η^2^ = 0.124; *M* = 7.53 cm, *SD* = 0.29 for low reward vs. *M* = 8.6, *SD* = 0.29 for high reward). For experienced mind wandering, the participants reported higher values for low compared to high reward trials (*t*(24) = 2.61, p = .016, η^2^ = 0.061; *M* = 5.52 cm, *SD* = 0.5 for low reward vs. *M* = 4.61, *SD* = 0.5 for high reward). One participant failed to complete the post-experimental questionnaire and did not report values for mind wandering.

### Time-frequency analysis on sensor-level data

The analysis of the time-frequency decomposed EEG data at sensor-level using cluster-based permutation tests revealed two significant clusters for the factor reward, two significant clusters for the factor task-switch, and no significant factor for the interaction. These clusters will be referred to as Cluster 1 to 4. The location of the observed clusters in time × frequency × sensor space, the corresponding effect sizes, as well as the condition-specific event-related spectral perturbations (ERSP) are illustrated in Figure 3. Frequency band specific power for each condition for frontal and posterior recording sites is shown in Figure 4.

**Figure 3:**
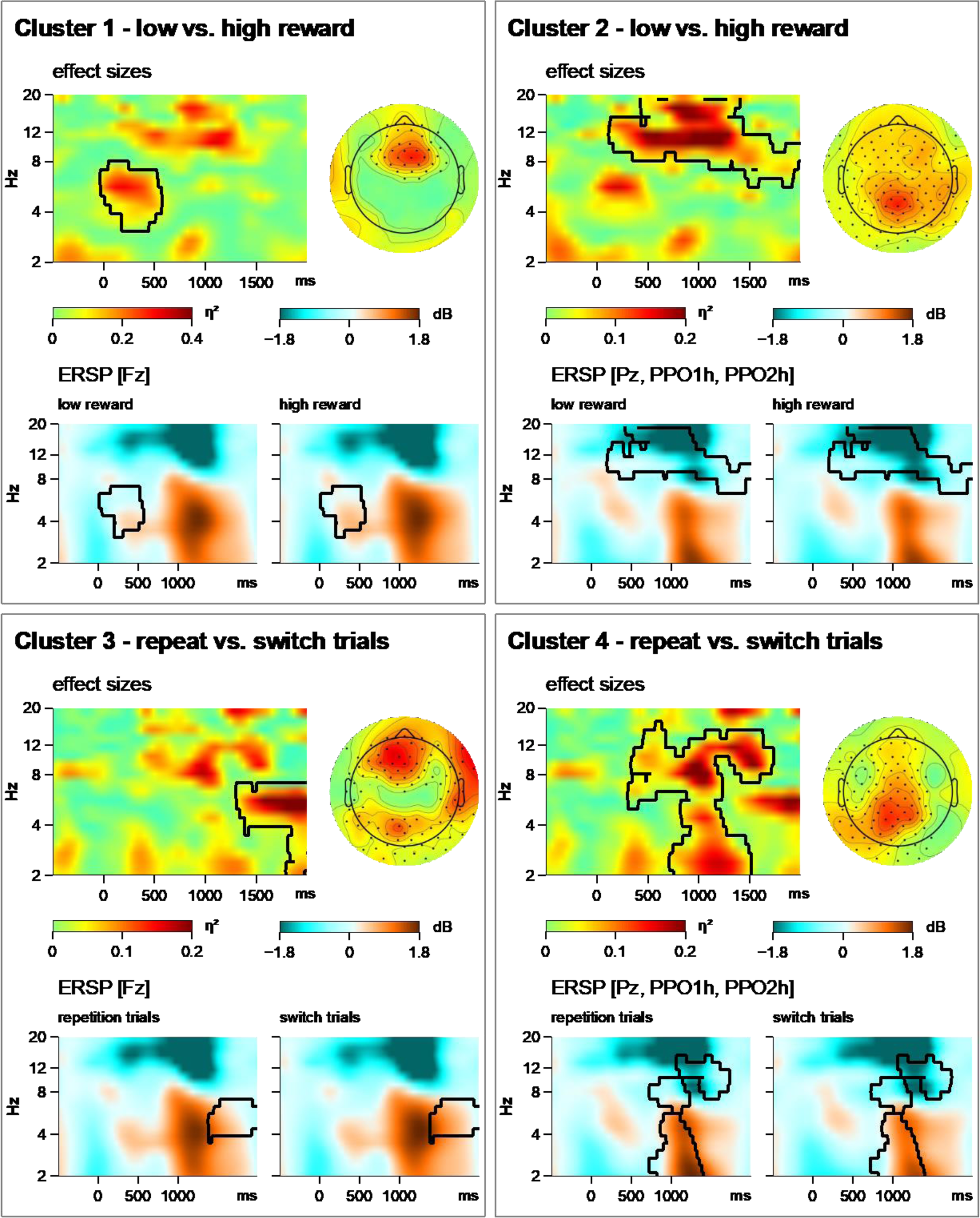
This figure depicts the result from the cluster-based permutation test performed on time-frequency decomposed sensor-level data. The onset of the task-cue is at 0 ms, the target display is presented at 800 ms. Four significant clusters were identified. For each cluster, effect sizes are displayed in time × frequency space (averaged across significant sensors) and as a topography (averaged across significant time-points and frequencies). Also, for each cluster, spectral power in time × frequency space is depicted for each condition for selected sensors.

**Figure 4:**
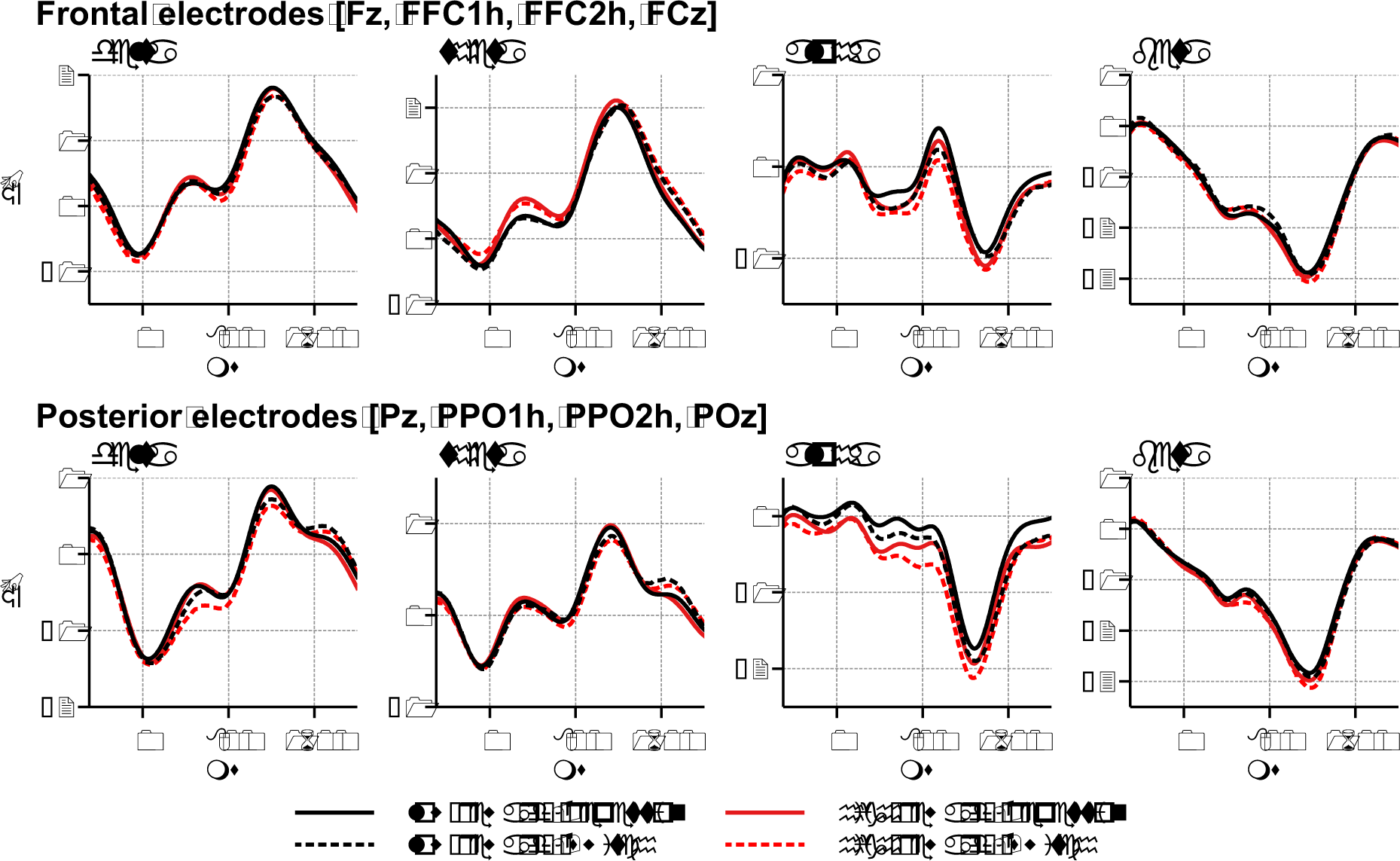
This figure shows the ERSPs for all conditions averaged for the delta (<4 Hz), theta (4 to 7 Hz), alpha (8 to 12 Hz), and beta (14 to 20 Hz) bands. Frequency band specific ERSPs are depicted as line plots for frontal and posterior electrodes. The onset of the task-cue is at 0 ms, the target display is presented at 800 ms.

Cluster 1 indicates a significant difference between the low and the high reward condition with low to medium effect sizes (c.f. J. Cohen, 1992). It is located in the theta-range during the cue-target interval at fronto-central electrodes. Mid-frontal theta-power during the cue-target interval is significantly higher in high versus low reward trials.

Cluster 2 also shows a significant difference between low and high reward trials with low to medium effect sizes. It is significant at almost all electrodes, but the largest effect sizes can be observed at mid-posterior electrodes. In time, Cluster 2 begins after the task-cue and spans over the whole trial (cue-target interval, probe display, response time-window). The largest effects can be observed in the alpha band and the lower beta band. It suggests a stronger alpha desynchronization, particularly at posterior recording sites, in high compared to low reward trials.

Cluster 3 suggests a significant effect of the factor task-switch with small to medium effect sizes. It is in the theta range after the response time window. The topography of effect sizes does not exhibit a clear focus, but the largest effect sizes can be observed at fronto-central electrodes. Here, theta power is stronger in switch-compared to repeat-trials.

Cluster 4 indicates a significant effect of task-switching, again with small to medium effect sizes. It is significant at almost all electrodes with the largest effect sizes at posterior sensors. It is significant in the delta, theta, and alpha range after the presentation of the probe display. Effects in the alpha band begin already during the cue-target interval and span the response time window as well. Event related spectral power in Cluster 4 is higher in repeat-compared to switch-trials.

### Source-reconstruction

Based on the results from the sensor-level analysis, four source-level analyses were conducted: For Cluster 1, low and high reward trials were compared for the time range from 0 to 600 ms at a frequency of 6 Hz (with ±2 Hz spectral smoothing). For Cluster 2, low and high reward trials were compared for the time range of 300 to 1300 ms at a frequency of 12 Hz (with ±2 Hz spectral smoothing). For Cluster 3, switch and repeat trials were compared for the time range from 1500 to 2200 ms at a frequency of 5.5 Hz (with ±2 Hz spectral smoothing). For Cluster 4, switch and repeat trials were compared for the time range of 700 to 1200 ms at a frequency of 9 Hz (with ±2 Hz spectral smoothing). To give an overview, the results of the source-level analyses have been projected on a cortical sheet. The resulting cortical sheet plots are depicted in Figure 5. The results listed below give an overview of the brain areas related to the observed effects. A detailed list of the anatomical labels for the voxels in significant clusters is provided in the supplementary material.

**Figure 5:**
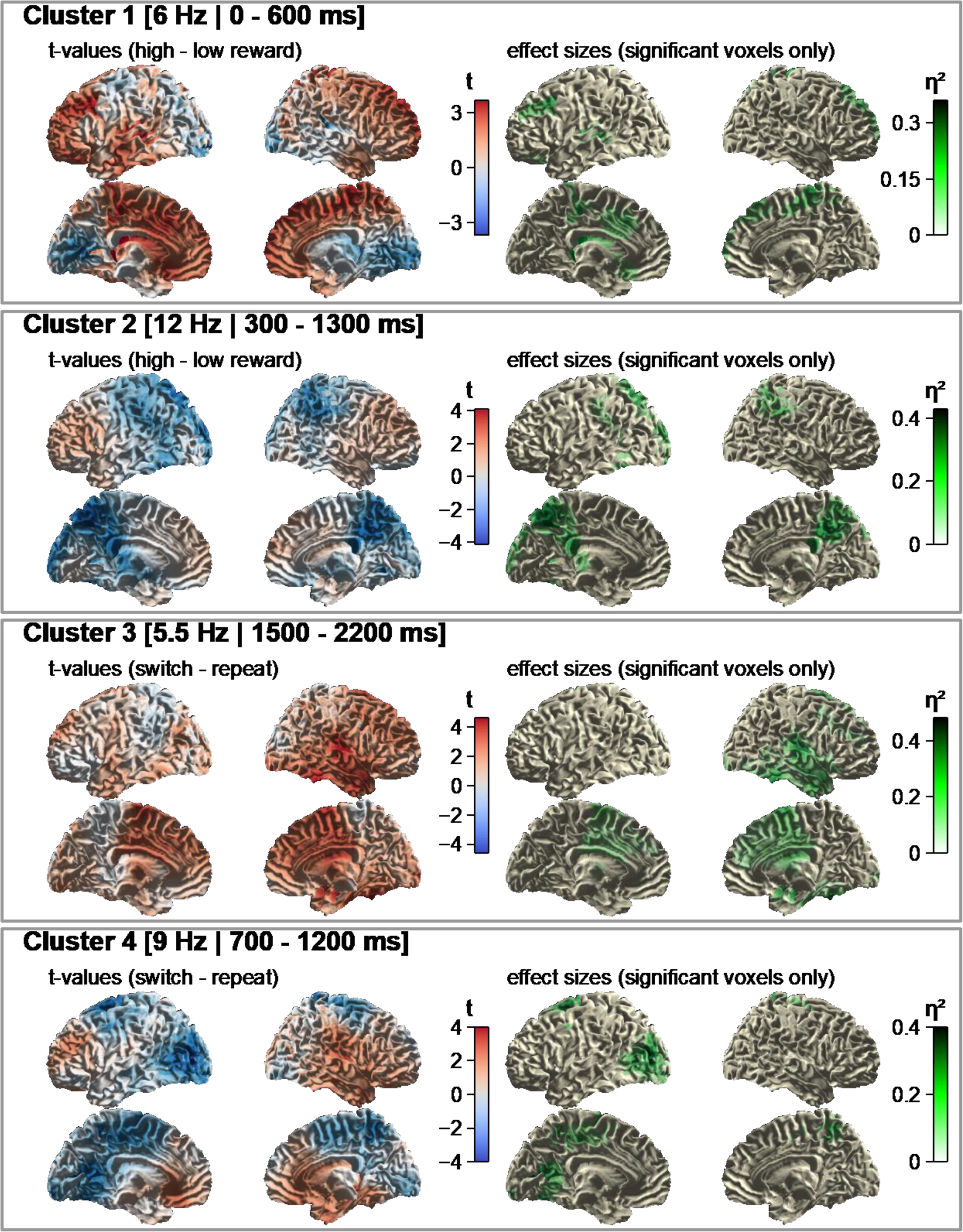
For each cluster, this figure shows the mass-univariate test-statistics (*t*-values) for all point-to-point comparisons in the source-space (left side). After controlling for multiple-comparisons using cluster-based permutation testing, the significant clusters are depicted by highlight the effect-sizes for all voxels belonging to a significant cluster.

The source-level analysis for Cluster 1 localizes the sources for the difference between low and high reward trials in the theta band during the cue-target interval. The results indicate that theta power was significantly higher for high versus low reward trials in frontal areas (superior, middle, and inferior frontal gyri and medial frontal cortex), motor areas (precentral gyri, supplementary motor area, paracentral lobules), left temporal areas, parietal regions (postcentral gyri, superior parietal lobules, precuneus), anterior and mid-cingulate cortex, as well as in left sub-cortical areas (caudate, putamen, thalamus).

For Cluster 2, the source level analysis investigates alpha power in a time-window covering most of the trial. Alpha power was higher in low compared to high reward trials in parietal areas (superior and inferior parietal lobules, postcentral gyri, paracentral lobules, precuneus, angular gyri, supra marginal gyri), occipital areas (superior, middle, and inferior occipital gyri, cuneus, lingual gyri), left temporal areas, the cerebellum, as well as in the mid- and posterior-cingulate cortex.

Cluster 3 is in the theta range in a time-window after the response. The source-level analysis revealed a higher theta power in switch versus repeat trials in right frontal regions (superior, middle, inferior gyrus), left and right superior medial frontal gyri, right parietal areas (superior and inferior lobules, precuneus, angular gyrus, supra-marginal gyrus), motoric areas (precentral gyrus, supplementary motor area, left paracentral lobule), right temporal areas, the right occipital cortex (superior, middle, and inferior occipital gyri, cuneus, lingual gyri), sub-cortical areas (caudate, putamen, thalamus), and the cerebellum.

The source-level analysis for Cluster 4 localizes the sources of alpha power after the presentation of the probe display. The results show increased alpha power in repeat compared to switch trials in frontal regions (superior and middle frontal gyri), motor areas (precentral gyri, supplementary motor area, paracentral lobules), parietal areas (superior parietal lobules, postcentral gyri, precuneus), mid- and posterior-cingulate cortex, left occipital areas (superior, middle, and inferior occipital gyri, cuneus, calcarine gyrus), and the cerebellum.

## Discussion

The present study investigated reward-related modulations of cognitive effort in a cued switching-task by means of the EEG. Besides behavioral measures and subjective ratings, the EEG was analyzed using time-frequency analysis. Subsequently, the effects observed on sensor level were localized in source-space using a beamforming approach. The behavioral data clearly indicates that the manipulation of cognitive effort was successful. The participants responded significantly faster in high versus low reward trials. The responses were also descriptively more accurate for high reward trials, but this difference was not significant. In addition, the participants reported significantly higher levels of motivation and effort for high reward trials, as well as significantly lower levels of mind wandering, in a post-experimental questionnaire. Responses were also significantly faster and more accurate in repeat compared to switch trials. This was expected, as the necessary updating of the task set in switch trials impedes performance (Kiesel et al., 2010). In line with previous studies (Capa et al., 2013; Hall-McMaster et al., 2019), no interaction of the factors reward and task-switch could be observed. This indicates that the prospect of reward boosted performance on a general level, irrespective of the specific control demand of the respective trial.

The analysis of time-frequency decomposed sensor-level EEG revealed two significant clusters for the factor reward-condition. Cluster 1 is in the theta frequency-range during the cue-target interval. The largest effect-sizes can be observed at fronto-central recording sites. This finding indicates that frontal midline theta power was significantly stronger in high versus low reward trials. Event-related frontal midline theta is a well-known correlate of the exertion of cognitive control (Cavanagh & Frank, 2014; Cavanagh & Shackman, 2015; Helfrich & Knight, 2016). Although no recent study reported an increased frontal midline theta related to a reward-related manipulation of cognitive effort, there is evidence that this connection might exists. Modulations of cognitive control caused by factors related to task-demands or motivation are reflected in midfrontal theta activity. For example, frontal theta has been found to be increased in conflict processing (Nigbur et al., 2011), as well as by high task demands (Fairclough & Ewing, 2017; Onton et al., 2005). It has also been observed to be decreased when motivation declines with time on task (Arnau et al., 2021). The experiment was also designed in a way to allow for disentangling top-down cognitive control in response to the task cue from target processing and response preparation. The increased frontal midline theta power during the cue-target interval reflected in Cluster 1 is therefore likely to reflect an increase in proactive control linked to an increase in mental effort (Braver, 2012; Braver et al., 2008). A reward-contingent increase of theta power during preparatory time-periods has also been observed in a recent study on rewarded task-prioritization, where the rewards associated with two different tasks were manipulated on a trial-to-trial basis (Liegel et al., 2022).

The results from the source localization support this interpretation. Theta activity was increased in high reward trials in the frontal and parietal regions, as well as in the cingulate cortex. These regions have been linked to the exertion of cognitive control and have been found to be sensitive to reward manipulations in fMRI studies (Egner, 2017; Jimura et al., 2010; Krebs et al., 2012; Padmala & Pessoa, 2011). It is hypothesized that cognitive control is established by orchestrating task-relevant brain areas using inter-areal communication in the theta range (M. M. Botvinick, 2007; Helfrich & Knight, 2016; Hopfinger et al., 2000). The cingulate cortex in particular has been described as being responsible to integrate reward-related information and cognitive control to adapt attentional processes, strategies, and behavior (Chong et al., 2017; Parro et al., 2018; Shenhav et al., 2013, 2016). On source level, the increase in proactive control might also be reflected by the increased theta power in temporal areas, related to attentional selection (Ramezanpour & Fallah, 2022), and in motor areas, possibly related to the preparation of motor-plans according to the task-rules (c.f. Pellegrino et al., 2018).

Another effect of the manipulation of effort is reflected in Cluster 2. It shows that the alpha power decrease throughout the trial is less pronounced in high versus low reward trials. The effect is strongest at centro-parietal electrodes. Alpha power has been described to reflect attentional biasing, with lower levels of alpha power being associated with an external focus of attention (c.f. Klimesch, 2012), and higher levels of alpha power indicating that attention is being directed towards mind-wandering, internal thoughts, and mental imagery (Arnau et al., 2020; Compton et al., 2019; Hanslmayr et al., 2011; Klimesch, 2012). In source-space, the strongest effect sizes can be observed at parieto-occipital areas and the posterior-cingulate cortex. These regions are associated with visual processing and action-preparation (Wang et al., 2019). Cluster 2 is therefore likely to reflect a stronger biasing of ex external (visual) attention over an internal focus of attention in high versus low reward trials.

The time-frequency analysis also revealed significant differences between repeat and switch trials. Cluster 3 is in the theta range in a time-window after the response with a frontocentral topography. This increase of frontal theta power after the response probably reflects an adjustment of control in preparation for the next trial. This could be triggered by conflict monitoring processes reacting to higher control demands in switch compared to repeat trials (Cavanagh & Frank, 2014; Nigbur et al., 2011). Another explanation is that the theta increase is error-related (c.f. Cavanagh & Frank, 2014), as the analysis of the EEG data also included trials with incorrect responses and the error rates were significantly higher in switch compared to repeat trials. Cluster 4 is in the delta, theta, and alpha range in a time window around the onset of the target display. The effect sizes are largest at centroparietal electrodes. The frequency-range as well as the mostly transient nature of most of the effect indicates that it reflects evoked activity in response to the onset of the target display. This activity is stronger for repetition than for switch trials suggesting a more pronounced phase coherence when no updating of the task rule is needed. On the other hand, the pre-stimulus alpha part of Cluster 4 shows a stronger alpha desynchronization for switch compared to repeat trials, which can again be interpreted as a stronger task orientation of attention (Klimesch, 2012). In line with this interpretation, the source-space analysis revealed that the largest effect sizes associated with pre-stimulus alpha can be observed at the visual cortex.

## Conclusion

The goal of the present study was to investigate how cognitive effort is reflected in the time-frequency space of the event-related EEG and to reconstruct the associated sources. The findings of the present study clearly show that the manipulation of cognitive effort using a monetary incentive manipulation (c.f. Knutson et al., 2000) was successful. Performance was significantly increased in high versus low reward trials and the participants also reported significantly higher motivation and cognitive effort for these trials. No interaction with the factor task-switching could be observed. This suggests that the experimental manipulation affected effort on a general level. This might be due to the fact that the reward was not directly dependent on response times, but was fully obtained if responses were simply fast enough (c.f. Frömer et al., 2021). Given that the accuracy was still high in switch-trials, there potentially was no need for the participants to emphasize effort in switch trials specifically. As a limitation of the study, it might also be the case that the sample size was too small to detect potential smaller effects. In the EEG data, the increase of cognitive effort was reflected by an increased mid-frontal theta activity during the cue-target interval, as well as by a more pronounced alpha-desynchronization throughout the trial. This indicates that cognitive effort was implemented by an increase of proactive control (Braver, 2012) as well as by an increased bias of attention towards external stimuli. The reconstructed sources for these effects are in areas related to cognitive control, and visual processing, respectively.

## Code and data availability

The code used for preprocessing, cleaning, and analyzing the data presented in this study can be found at https://github.com/stefanarnau/theta_cognitive_effort. The preprocessed EEG data, the head-models, source-models, and individual electrode positions can be obtained from https://osf.io/ndgst/.

## Author Contributions

**Stefan Arnau:** Conceptualization, Formal Analysis, Methodology, Visualization, Writing – Original Draft. **Nathalie Liegel:** Formal Analysis, Methodology, Visualization, Writing – Review & Editing. **Edmund Wascher:** Conceptualization, Writing – Review & Editing.

## Acknowledgements

We would like to thank Erhan Genç, Christoph Fraenz, and Michael Burke for setting up and organizing the acquisition of the structural MRI scans. We would also like to thank Pia Deltenre, and Barbara Foschi for their assistance in collecting the data, as well as Tobias Blanke for programming the experiment.

