## Supplementary material for "Frontal midline theta power during the cue-target-interval reflects increased cognitive effort in rewarded task-switching"

This document lists the labels of brain regions associated with voxels of significant clusters.

**Cluster 1**

{'Precentral_L' }
{'Precentral_R' }
{'Frontal_Sup_L' }
{'Frontal_Sup_R' }
{'Frontal_Sup_Orb_L' }
{'Frontal_Sup_Orb_R' }
{'Frontal_Mid_L' }
{'Frontal_Mid_R' }
{'Frontal_Mid_Orb_L' }
{'Frontal_Inf_Oper_L' }
{'Frontal_Inf_Tri_L' }
{'Frontal_Inf_Orb_L' }
{'Rolandic_Oper_L' }
{'Supp_Motor_Area_L' }
{'Supp_Motor_Area_R' }
{'Olfactory_L' }
{'Frontal_Sup_Medial_L'}
{'Frontal_Sup_Medial_R'}
{'Frontal_Med_Orb_L' }
{'Frontal_Med_Orb_R' }
{'Rectus_L' }
{'Rectus_R' }
{'Insula_L' }
{'Cingulum_Ant_L' }
{'Cingulum_Ant_R' }
{'Cingulum_Mid_L' }
{'Cingulum_Mid_R' }
{'Cingulum_Post_L' }
{'Hippocampus_L' }
{'ParaHippocampal_L' }
{'Cuneus_R' }
{'Lingual_L' }
{'Occipital_Sup_R' }
{'Fusiform_L' }
{'Postcentral_L' }
{'Postcentral_R' }
{'Parietal_Sup_L' }
{'Parietal_Sup_R' }
{'Precuneus_L' }
{'Precuneus_R' }
{'Paracentral_Lobule_L'}
{'Paracentral_Lobule_R'}
{'Caudate_L' }

{'Putamen_L' }
{'Pallidum_L' }
{'Thalamus_L' }
{'Heschl_L' }
{'Temporal_Sup_L' }
{'Temporal_Pole_Sup_L' }
{'Temporal_Mid_L' }
{'Temporal_Inf_L' }

**Cluster 2:**

{'Precentral_L' }
{'Precentral_R' }
{'Frontal_Mid_R' }
{'Cingulum_Mid_L' ????
{'Cingulum_Mid_R' ???
{'Cingulum_Post_L' }
{'Cingulum_Post_R' }
{'Hippocampus_L' }
{'Calcarine_L' }
{'Cuneus_L' }
{'Cuneus_R' }
{'Lingual_L' }
{'Lingual_R' }
{'Occipital_Sup_L' }
{'Occipital_Sup_R' }
{'Occipital_Mid_L' }
{'Occipital_Mid_R' }
{'Occipital_Inf_L' }
{'Fusiform_L' }
{'Postcentral_L' }
{'Postcentral_R' }
{'Parietal_Sup_L' }
{'Parietal_Sup_R' }
{'Parietal_Inf_L' }
{'Parietal_Inf_R' }
{'SupraMarginal_L' }
{'SupraMarginal_R' }
{'Angular_L' }
{'Angular_R' }
{'Precuneus_L' }
{'Precuneus_R' }
{'Paracentral_Lobule_L'}
{'Paracentral_Lobule_R'}
{'Thalamus_L' }
{'Thalamus_R' }
{'Temporal_Sup_L' }

{'Temporal_Mid_L' }
{'Temporal_Inf_L' }
{'Cerebellum_Crus1_L' }
{'Cerebellum_Crus2_L' }
{'Cerebellum_6_L' }
{'Cerebellum_7b_L' }
{'Cerebellum_8_L' }

**Cluster 3:**

{'Precentral_L' }
{'Precentral_R' }
{'Frontal_Sup_L' }
{'Frontal_Sup_R' }
{'Frontal_Mid_R' }
{'Frontal_Inf_Oper_R' }
{'Frontal_Inf_Tri_R' }
{'Frontal_Inf_Orb_R' }
{'Rolandic_Oper_R' }
{'Supp_Motor_Area_L' }
{'Supp_Motor_Area_R' }
{'Frontal_Sup_Medial_L'}
{'Frontal_Sup_Medial_R'}
{'Insula_R' }
{'Cingulum_Ant_L' }
{'Cingulum_Ant_R' }
{'Cingulum_Mid_L' }
{'Cingulum_Mid_R' }
{'Cingulum_Post_R' }
{'Hippocampus_R' }
{'ParaHippocampal_R' }
{'Amygdala_R' }
{'Calcarine_R' }
{'Cuneus_R' }
{'Lingual_L' }
{'Lingual_R' }
{'Occipital_Sup_R' }
{'Occipital_Mid_R' }
{'Occipital_Inf_R' }
{'Fusiform_R' }
{'Postcentral_R' }
{'Parietal_Sup_R' }
{'Parietal_Inf_R' }
{'SupraMarginal_R' }
{'Angular_R' }
{'Precuneus_R' }
{'Paracentral_Lobule_L'}

{'Caudate_L' }
{'Caudate_R' }
{'Putamen_R' }
{'Pallidum_R' }
{'Thalamus_L' }
{'Thalamus_R' }
{'Heschl_R' }
{'Temporal_Sup_R' }
{'Temporal_Pole_Sup_R' }
{'Temporal_Mid_R' }
{'Temporal_Pole_Mid_R' }
{'Temporal_Inf_R' }
{'Cerebellum_Crus1_L' }
{'Cerebellum_Crus1_R' }
{'Cerebellum_Crus2_L' }
{'Cerebellum_Crus2_R' }
{'Cerebellum_3_R' }
{'Cerebellum_4_5_R' }
{'Cerebellum_6_R' }
{'Cerebellum_7b_L' }
{'Cerebellum_7b_R' }
{'Cerebellum_8_L' }
{'Cerebellum_8_R' }
{'Cerebellum_9_R' }
{'Cerebellum_10_R' }
{'Vermis_8' }
{'Vermis_9' }

**Cluster 4:**

{'Precentral_L' }
{'Precentral_R' }
{'Frontal_Sup_L' }
{'Frontal_Sup_R' }
{'Frontal_Mid_L' }
{'Frontal_Mid_R' }
{'Supp_Motor_Area_L' }
{'Supp_Motor_Area_R' }
{'Cingulum_Mid_L' }
{'Cingulum_Mid_R' }
{'Cingulum_Post_L' }
{'Cingulum_Post_R' }
{'Calcarine_L' }
{'Calcarine_R' }
{'Cuneus_L' }
{'Lingual_L' }
{'Occipital_Sup_L' }

```
{'Occipital_Mid_L'     }
{'Occipital_Inf_L'     }
{'Fusiform_L'          }
{'Postcentral_L'       }
{'Postcentral_R'       }
{'Parietal_Sup_L'      }
{'Parietal_Sup_R'      }
{'Angular_L'           }
{'Precuneus_L'         }
{'Precuneus_R'         }
{'Paracentral_Lobule_L'}
{'Paracentral_Lobule_R'}
{'Temporal_Mid_L'      }
{'Temporal_Inf_L'      }
{'Cerebellum_Crus1_L'  }
{'Cerebellum_Crus2_L'  }
{'Cerebellum_4_5_L'    }
{'Cerebellum_6_L'      }
{'Vermis_4_5'          }
```
